# Increased deformations are dispensable for cell mechanoresponse in engineered bone analogs mimicking aging bone marrow

**DOI:** 10.1101/2023.09.24.559187

**Authors:** Alexander M Regner, Maximilien DeLeon, Kalin D. Gibbons, Sean Howard, Derek Q. Nesbitt, Trevor J. Lujan, Clare K. Fitzpatrick, Mary C Farach-Carson, Danielle Wu, Gunes Uzer

## Abstract

Aged individuals and astronauts experience bone loss despite rigorous physical activity. Bone mechanoresponse is in-part regulated by mesenchymal stem cells (MSCs) that respond to mechanical stimuli. Direct delivery of low intensity vibration (LIV) recovers MSC proliferation in senescence and simulated microgravity models, indicating that age-related reductions in mechanical signal delivery within bone marrow may contribute to declining bone mechanoresponse. To answer this question, we developed a 3D bone marrow analog that controls trabecular geometry, marrow mechanics and external stimuli. Validated finite element (FE) models were developed to quantify strain environment within hydrogels during LIV. Bone marrow analogs with gyroid-based trabeculae of bone volume fractions (BV/TV) corresponding to adult (25%) and aged (13%) mice were printed using polylactic acid (PLA). MSCs encapsulated in migration-permissive hydrogels within printed trabeculae showed robust cell populations on both PLA surface and hydrogel within a week. Following 14 days of LIV treatment (1g, 100 Hz, 1 hour/day), type-I collagen and F-actin were quantified for the cells in the hydrogel fraction. While LIV increased all measured outcomes, FE models predicted higher von Mises strains for the 13% BV/TV groups (0.2%) when compared to the 25% BV/TV group (0.1%). Despite increased strains, collagen-I and F-actin measures remained lower in the 13% BV/TV groups when compared to 25% BV/TV counterparts, indicating that cell response to LIV does not depend on hydrogel strains and that bone volume fraction (i.e. available bone surface) directly affects cell behavior in the hydrogel phase independent of the external stimuli. Overall, bone marrow analogs offer a robust and repeatable platform to study bone mechanobiology.

## 1. Introduction

Mesenchymal stem cells (MSC) found in adult musculoskeletal tissues maintain tissue turnover and repair. In bone, MSC is the common progenitor for osteoblasts, osteocytes and adipocytes.[1–3] During habitual loading, MSCs provide necessary osteoblast populations to facilitate bone modeling.[4] Aging[5–7] and chronic unloading under microgravity[8, 9] decrease exercise-induced bone accrual. While aging and unloading phenotypes are associated with decreased proliferative and osteogenic differentiation capacity of MSCs,[10, 11] findings from *in vitro* replicative senescence[12] and simulated microgravity models[13] show that exposing MSCs to mechanically active environments improve these outcomes. *In vivo* studies using long daily treadmill activity[14] or ladder climbing regimens[15] report improved MSC function with long-term physical activity. These studies suggest that trabecular volume may contribute to declining bone mechanoresponse in aging and prolonged unloading scenarios. To this end, we sought to develop a 3D bone marrow analog to study the role of trabecular bone volume on MSC function and quantify the effects of trabecular bone volume on MSC mechanoresponse under external mechanical stimulation.

MSCs in bone exist in a mechanically rich environment. Inside the bone marrow, bone surfaces where bone cells reside, are exposed to matrix deformations[16–19], accelerations[20–25], fluid flow[26–30], and changes in intramedullary pressure[31–33], all of which are inseparable[34]. Cyclic bone deformations associated with walking, for example, generates strains up to 400µɛ and fluid velocities up to 100 µm/s within the lacunar– canalicular network [35]. When looking for candidate loading regimens for the bone marrow analog model, we determined that vibration is the most applicable and practical mechanical input. When a person runs, tibial accelerations reach 2–5g[36] (1g = 9.81 m/s^2^), creating complex loading conditions[37] capable of driving MSC function.[38] High frequency and low magnitude signals are part of bone physiology; measuring 24 hour strain history of load bearing bones across different species showed that bones are subjected to thousands of very small, high frequency strains (< 10 µɛ) during the day.[39] Leveraging the presence of these small strains, application of low intensity vibration (LIV), usually applied between 30 and 100Hz with acceleration magnitudes of 0.1 to 1g, have been used in clinical, pre-clinical and cellular studies[4]. In clinical studies, LIV protects bone quantity and quality in women with osteoporosis[40, 41], children with cerebral palsy[42], and improves bone indices in child cancer survivors[43]. Animal studies demonstrated that external LIV application increases trabecular bone density and volume[20], creates stiffer bones [44], and improves disuse-induced bone loss[45]. At the cellular level, our group reported that application of LIV increases MSC contractility [46], activates RhoA signaling [47], and results in increased osteogenic differentiation and proliferation of MSCs[48]. Similar to other mechanical signals such as substrate strain, LIV increases the nuclear accumulation of mechanically-sensitive transcription factors, β-catenin[49] and YAP[50] as well as increasing nuclear stiffness[51]. We recently reported that in a simulated microgravity model where MSCs were exposed to rotating cell culture vessels for 72h to simulate gravitational unloading, LIV rescued YAP nuclear entry induced by RhoA-activator lysophosphatidic acid[50].

The mechanical environment LIV generates within the bone marrow is not well studied due to a lack of repeatable model systems. Early modeling efforts utilizing idealized rectangular bone lattices[37] or realistic bone geometries from µCT scans of a human lumbar vertebra[52] reported that LIV can generate fluid shear stress on bone surfaces up to 1-2 Pa depending on the bone marrow viscosity. Follow-up *ex vivo* studies that combined computational modeling with trabecular explant models under LIV showed that LIV-induced fluid shear was, in part, correlated with histological findings. However, fluid shear only groups remained lower than the LIV groups[53], suggesting that other mechanical or geometrical factor plays a role. Supporting these findings, our group further reported that LIV effect was largely independent of fluid shear in MSCs[48], osteoblasts[54] and osteocytes[55]. While previous studies reveal possible LIV mechanisms, there are no current *in vitro* models to capture the mechanical complexity of bone marrow during mechanical stimulation. Consequently, biological variation and sample availability makes it challenging to setup robust *ex vivo models* to study the effect of trabecular bone volume using multiple outcomes from identical conditions.

Therefore, to overcome these challenges, we developed a cell-laden bone marrow analog with a 3D printed PLA trabeculae and hyaluronic acid-based bone marrow to quantify the effects of trabecular volume on MSC response during LIV. The bone marrow strains created during 1g, 100Hz vibrations in both 13% and 25% trabecular bone volumes – representing bone volumes of 64 week old (aged) and 8 week old (young) adult male C57BL/6J mouse – were compared[56]. Both validated finite element (FE) simulations and *in vitro* experiments were used to determine whether trabecular bone volume affects the bone marrow mechanical environment and MSC response.

### Experimental design

To create a validated experimental bone-analog model, PLA was chosen for the material of the trabecular structure due to its biocompatibility[57] and ease of use in 3D printing applications[58]. Previously developed hyaluronic acid-based hydrogels with MMP cleavable crosslinks and cell attachment sites were chosen to model bone marrow[59]. Primary mouse MSCs dispersed in hydrogels were used as the cell model due to their versatility in both proliferative and differentiation experiments utilized in previous studies[13, 16, 20, 44, 45, 47–50, 60]. To generate an experimentally validated FE model we first utilized a collagen-based gelatin hydrogel (i.e., calibration gels) followed by hyaluronic acid (HA) hydrogel experiments to set model parameters. To define hydrogel material properties for the FE model calibration we employed high-speed compression tests and high-speed speckle photography analysis. Due to large difference in elastic modulus between bone marrow and trabecular bone [57, 61, 62], PLA scaffolds were assumed to be rigid for the purposes of the FE model. Following FE model calibration, idealized 3D trabecular structures based on gyroid model[63] with either 13% or 25% volume fill (i.e. trabecular volume) were generated and LIV-induced HA hydrogel strains were compared between the two trabecular volumes (**Fig. 1a**). For *in vitro* experiments (**Fig. 1b**), we encapsulated 13% and 25% PLA scaffolds with HA hydrogel-MSC mixture using 1 million cells per mL of hydrogel. Cell-laden hydrogels with no PLA scaffold were used as no-scaffold controls (referred as 0%). Cell-laden 13% and 25% scaffolds were exposed to LIV vibration one hour every day at room temperature using 100 Hz frequency and 1g acceleration magnitude using protocols we established previously[48]. After 14 days, samples were allocated for mRNA (N=4/group), immunostaining (N=3/group). Controls were handled the same but were not vibrated.

**Figure 1.**
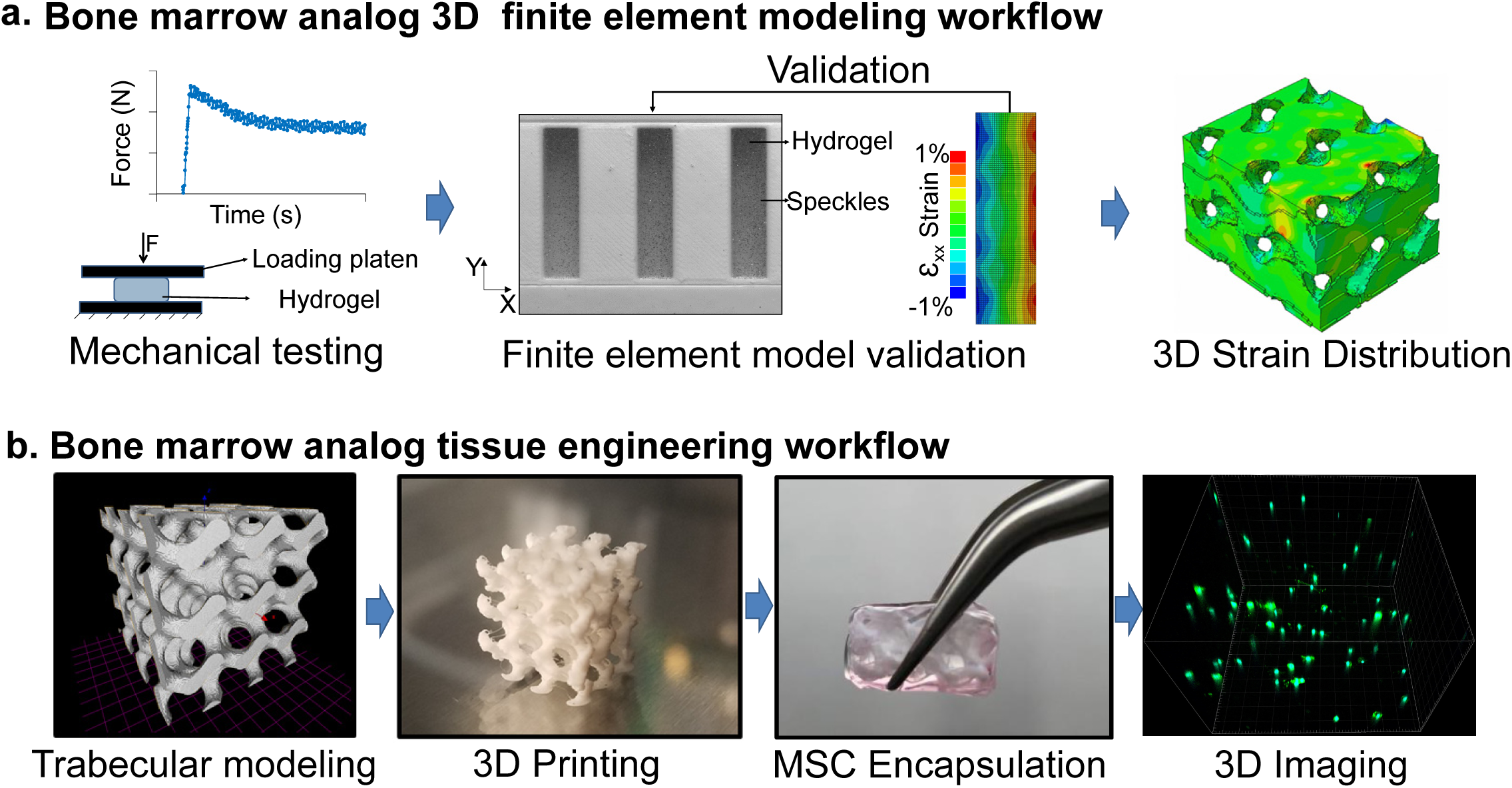
**Experimental Design**. **a)** Both calibration and HA gels were mechanically tested and Finite element (FE) models were experimentally calibrated to generate 3D strain distributions scaffolds with both 13% and 25% trabecular bone volumes – representing bone volumes of 64 week old (aged) and 8 week old (young) adult male C57BL/6J mouse. **b)** 3D bone analogs were generated using PLA printed trabeculae and primary MSCs were encapsulated using HA hydrogels with integrin binding motifs and metalloproteinases-sensitive crosslinkers. LIV was applied for 14 days (100Hz, 1g, 1hr/day) followed by volumetric imaging of scaffolds via confocal microscopy.

## 2. Methods and materials

### Preparation of hydrogels for mechanical testing and calibration

Calibration gels (Knox unflavored Gelatin) or HA hydrogels (Advanced Biomatrix: GS1004) were prepared in 3D PLA printed cylindrical molds with a diameter of 24 mm and a height of 9 mm (**Fig. 2a**). The calibration gels were prepared at 2 mg/ml via dissolving the gelatin powder in 50°C water. Sample molds used for calibration gels were lined with a thin layer of petroleum jelly to improve mold release and 4.1 ml of 2 mg/ml gelatin mixture was pipetted in. Filled molds were cooled over ice for 1 hour to cross-link and then both the gel and mold were deposited into a room temperature water bath to for 30 minutes. The calibration gel samples were carefully removed from the molds and tested. All tests were completed within 3 hours of initial calibration gel deposition into the molds. HA hydrogels were prepared according to manufacturer protocols. Briefly, degassed (DG) water was extracted via syringe and added to the Glycosil (5.0 ml DG water) and Extralink-Lite (2.5 ml DG water). These were vortexed on a rocker for 60 minutes and vigorously shaken by hand every 15 minutes. These solutions then were combined in a 4:1 ratio (v/v) and mixed by pipette into the mold. These gels cross-linked for 90 minutes at room temperature, removed from the mold, deposited into PBS in 90mm well plate (VWR#10062-878) sealed with parafilm, and left overnight in a 37°C in an incubator, and. The next day the HA hydrogels were tested one at a time, removing them from the PBS immediately prior to testing.

**Figure 2.**
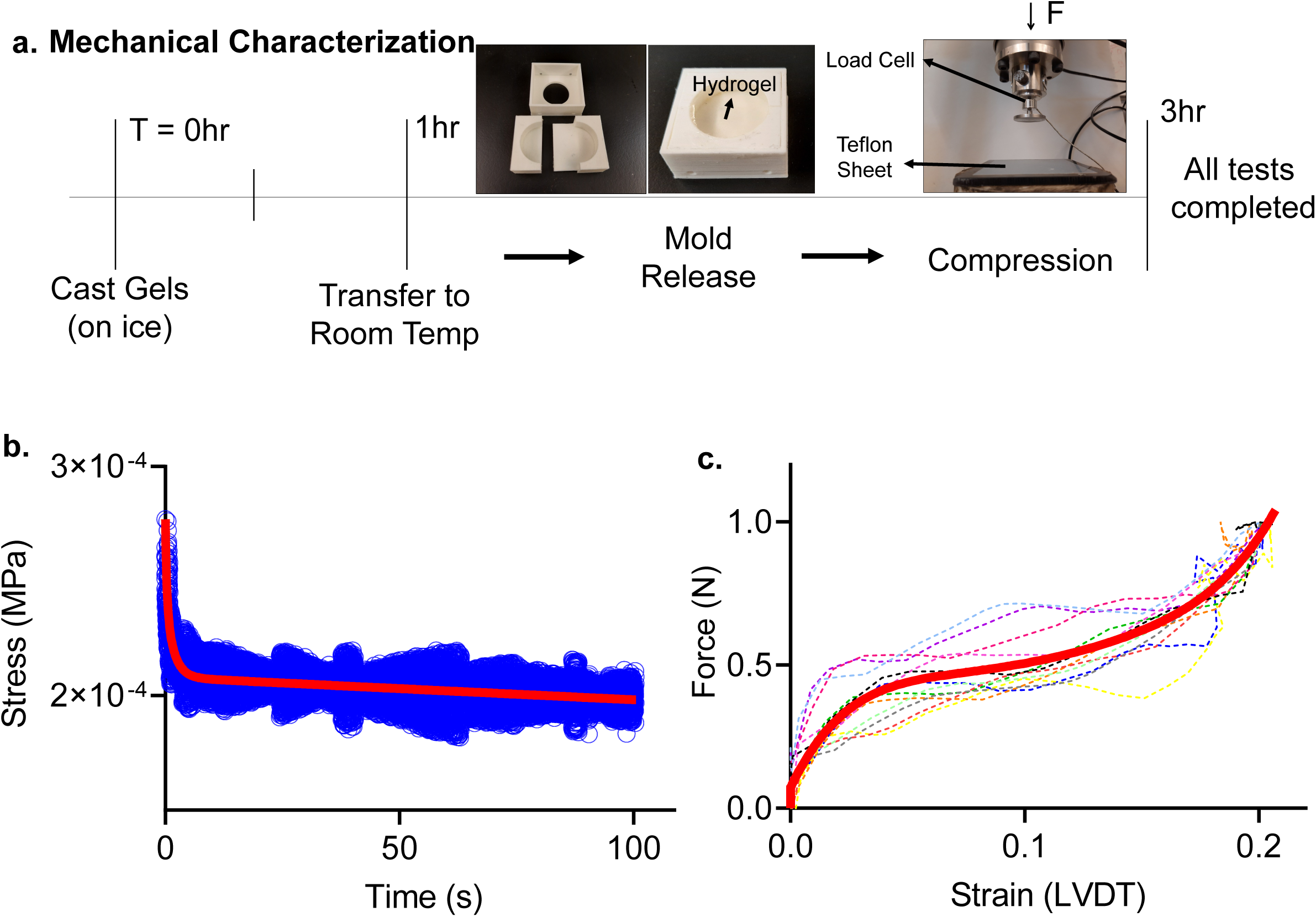
Mechanical Characterization Setup. a) Timeline of the compression testing, with all tests completed within 3 hours of initial gelation for the calibration gels. 3D printed mold used to cast instantaneous gel pucks (24mm internal diameter, and a height of 9mm). Experiments were completed using an INSTRON ElectroPuls E10000 using a 10N load-cell. The sample rested on a Teflon coated sheet lightly coated in water to prevent adherence. **b)** Relaxation data over a 100 second time period (blue) and the resulting fit line (red) relaxing from 2.8×10^-5^ MPa and reaching long term relaxation at approximately 2.1×10^-5^ Mpa. **c)** Instantaneous ramp data from 12 trials (dotted 5th order polynomial fit (blue), compressed to 0.2 strain (10.8 mm/s).

### Instantaneous compression testing

Instantaneous compression testing was performed using an INSTRON ElectroPuls E10000 using a 10N load-cell attached to a loading plate with a diameter of 35mm. At the start of each test, calibration and HA gels were placed on water-coated Teflon sheets to prevent adherence (**Fig. 2a**). Samples were pre-loaded with 0.025N to ensure full contact with the plate, then compressed to 20% strain (compared to pre-loaded gauge length) at a rate of 90 mm/s. Loading plate position then was held constant for 100 seconds and load-cell data was collected and exported into an excel sheet (**Fig. 2b**). The 100 second relaxation data was fitted using a 2-term exponential decay function [f(x) = a*e*^bx^ + c*e*^dx^] representing viscoelastic material relaxation.

Next, instantaneous force and displacement data was collected during 20% compression at a rate of 10.8 mm/s (**Fig. 2c**). Compression data was entered into MATLAB, converted to engineering stress and engineering strain, and then fit using a 5^th^ order polynomial [f(x) = ax^5^ + bx^4^ + cx^3^ + dx^2^ + ex +f] representing the instantaneous material behavior. Stress-strain curves for the calibration and HA gels are provided in Supplementary **Fig.S1**.

### Digital image correlation tests

Prepared calibration gels were tested in wells with differing width for FE model validation. Wells with fixed depth (9mm) and height (27.5mm) were 3D printed out of PLA for widths of 4mm, 5mm, 6mm, and 8mm (**Fig. 3a**). These wells were filled with calibration gels as described above and then were speckled with a 50/50 mixture of talc powder and grid 220 silicon carbide (SiC) to track their motions via speckle photometry as previously described^18^. To track light-reflective siC speckles, a white LED light source was used and the motion of the surface was captured with a Photron UX50 high-speed camera at a rate of 2000 frames per second (fps). PLA wells filled with calibration gels were vibrated horizontally at 100Hz with an acceleration of 1g via a custom-made vibration bioreactor driven by a Labworks ET-126HF-1,-4 (13lbf) Electrodynamic transducer using a sinusoidal driving function (**Fig. 3a**). All vibration tests were performed within 3h of gel casting. Recording of gel motion was started 15 seconds post-LIV to ensure that steady state was reached.

**Figure 3.**
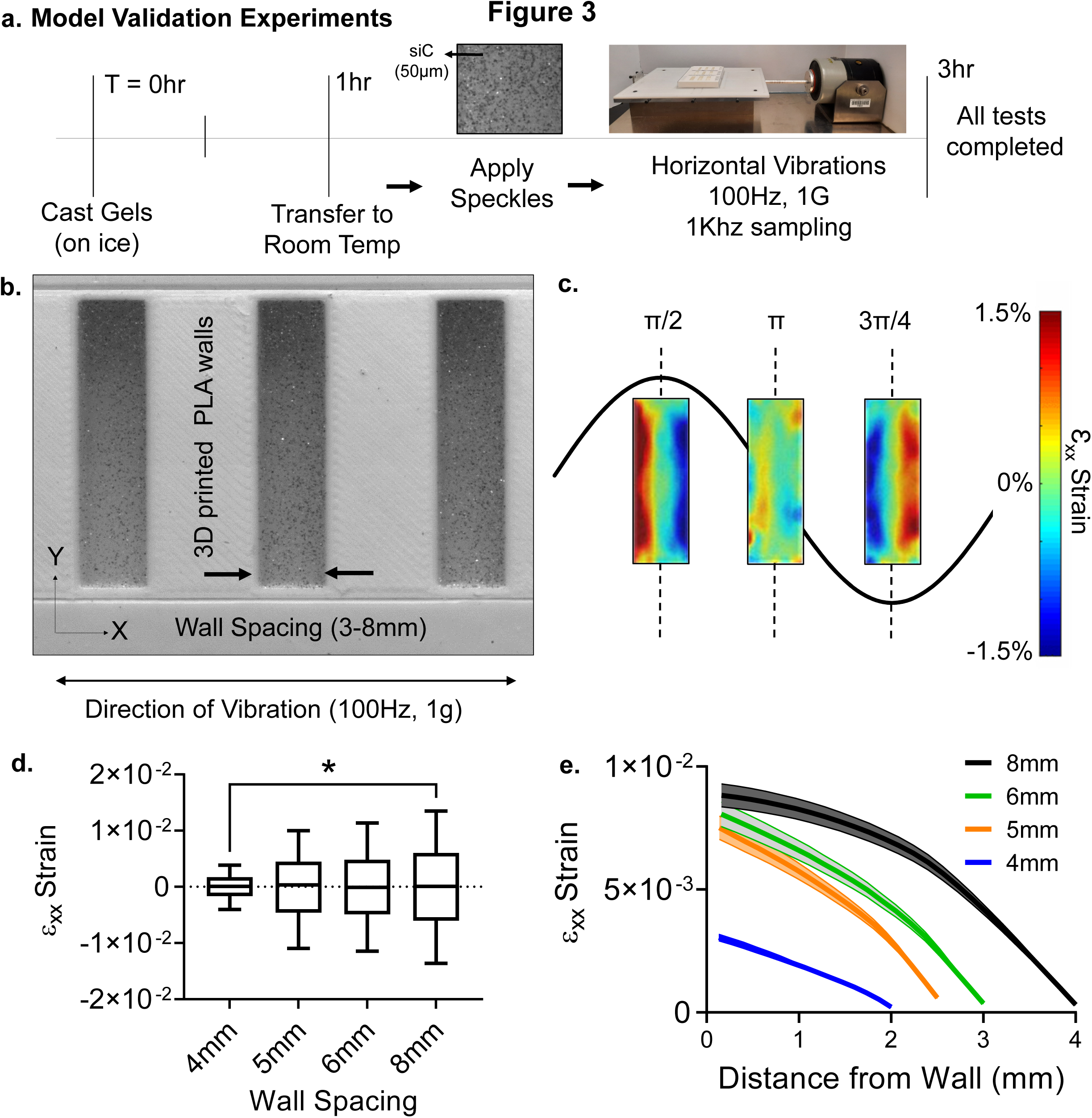
Model validation experiments. a) Timeline of the DIC tests, with all tests completed within 3 hours of initial gelation for the calibration gels. Hydrogels were speckled with 220 grid SiC and the camera recorded the motions at 2000 frames per second (fps). Horizontal vibration plate was driven by Labworks ET-126HF-1,-4 (13lbf) Electrodynamic transducer **b)** A PLA experimental setup was manufactured to visualize strain during100Hz –1g vibrations at the horizontal direction. ε_xx_ strains varied periodically with the maximum value of 0.015 at 8mm spacing. **c)** As the spacing decreased 62% to 4mm ε_xx_ strain decreased by two orders of magnitude showing a positive, non-linear correlation in the average maximum of the strain. **d)** Mapping max strain magnitudes from the wall to the mid-point showed a convex parabolic strain profile that minimized at the middle of the well.

Recorded high speed videos at 2000fps were analyzed using digital image correlation (DIC) with NCORR software[64] (V1.2) to export gel displacements relative to a moving frame of reference marked on the well surface. Post-processing of the full-field displacement maps were performed in MATLAB. Vibrations were applied along the x-axis and the well height direction was referred as y-axis (**Fig. 3b**). During the post-processing of the vibration experiments, the ε_xx_ strain was exclusively analyzed, as it was found to be at least one order of magnitude larger than ε_yy_ or ε_xy_ (see **Fig.S2**). For HA gel experiments, only the 8mm well width was used and same steps were repeated as above. All experiments were performed in triplicate.

### Finite element model generation and validation

A pre-processing software package (*Hypermesh v2017.2, Altair, MI)* was used to generate meshes for finite element analysis. 3D printed well geometries between 4 and 8mm were imported as STL files and converted to rigid R3D4 mixed elements. The hydrogels were meshed using hexahedral C3D8R elements with a size of 0.25 mm. These meshes were imported into a FE solver (*ABAQUS R2019×, Simula, RI)* to simulate vibration. The hydrogel material was modeled with a density of 1.0023 g/cm^3^ for calibration gels and a density of 1.0018 g/cm^3^ for HA gels. Hyper-viscoelastic material responses were modeled using the procedure outlined by Dalrymple^12^. For viscoelastic modeling, the long-term relaxation was measured starting from the end of the instantaneous compression to a 100 second interval. This then was fit using a second order exponential function, and data was extrapolated back to 10^-4^ seconds to ensure that input data to the finite element simulation was within the rate of vibration (100 Hz). For viscoelastic material definition a single-term Ogden model with three-term Prony Series was used[65]. The elastic material definition was modeled as hyperelastic with a 5^th^ order Ogden fit, 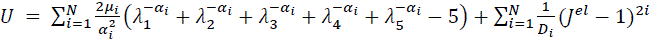 where λ_i_ are principal stretches, N is a parameter (in this case equal to 5), μ_i_, α_i_, and *D*_i_ are temperature dependent material parameters. These materials were defined within the FE solver using the instantaneous, uniaxial test data (**Fig. 1c**).

The material interaction between the undeformable and hydrogel parts used a linear press-overclosure with a slope of 1.0×10^-2^ and a friction and damping coefficients that minimized the difference between experimental and numerical ε_xx_ strain magnitudes at timepoint π/2 using a design of experiments optimization as described below. Mechanical vibration was applied using a connector actuator driven by a periodic velocity equation with an amplitude of 15.61 m/s, and a circular frequency of 100 Hz (628.31 radians)[54].

### Damping and friction coefficient optimization

The root mean square error (RMSE) between the time dependent FE displacement output and the DIC experimental data were calculated using the ε_xx_ surface strain at the π/2 timepoint of the 8mm well width. To calibrate the friction and damping coefficients in the FE model, a 3 factor 3 level design of experiments (DOE) was performed starting from the measured stiffness value. For the friction and damping coefficients, we started with ABAQUS recommended defaults values of 0.75 and 0.03, respectively. The stiffness value for the calibration gels varied ±50% of the measured value and the friction and damping coefficients varied between 0.1 –10 and 0.03-0.3, resulting in 27 simulations (**Table 1**). For each comparison, the gel surface was divided into a 15×37 grid and for each sub-region the RMSE difference between the experimental and computational ε_xx_ strains were compared. Following this analysis, the friction and damping coefficients that minimized the RMSE response were selected (**Fig. 4a**). Because increasing or decreasing stiffness from the measured values always resulted in larger RMSE values, 2 factor 3 level DOE was used for the HA gels (**Table S1**) and the friction and damping coefficients that minimized the RMSE response were selected (**Fig.S3a**).

**Figure 4.**
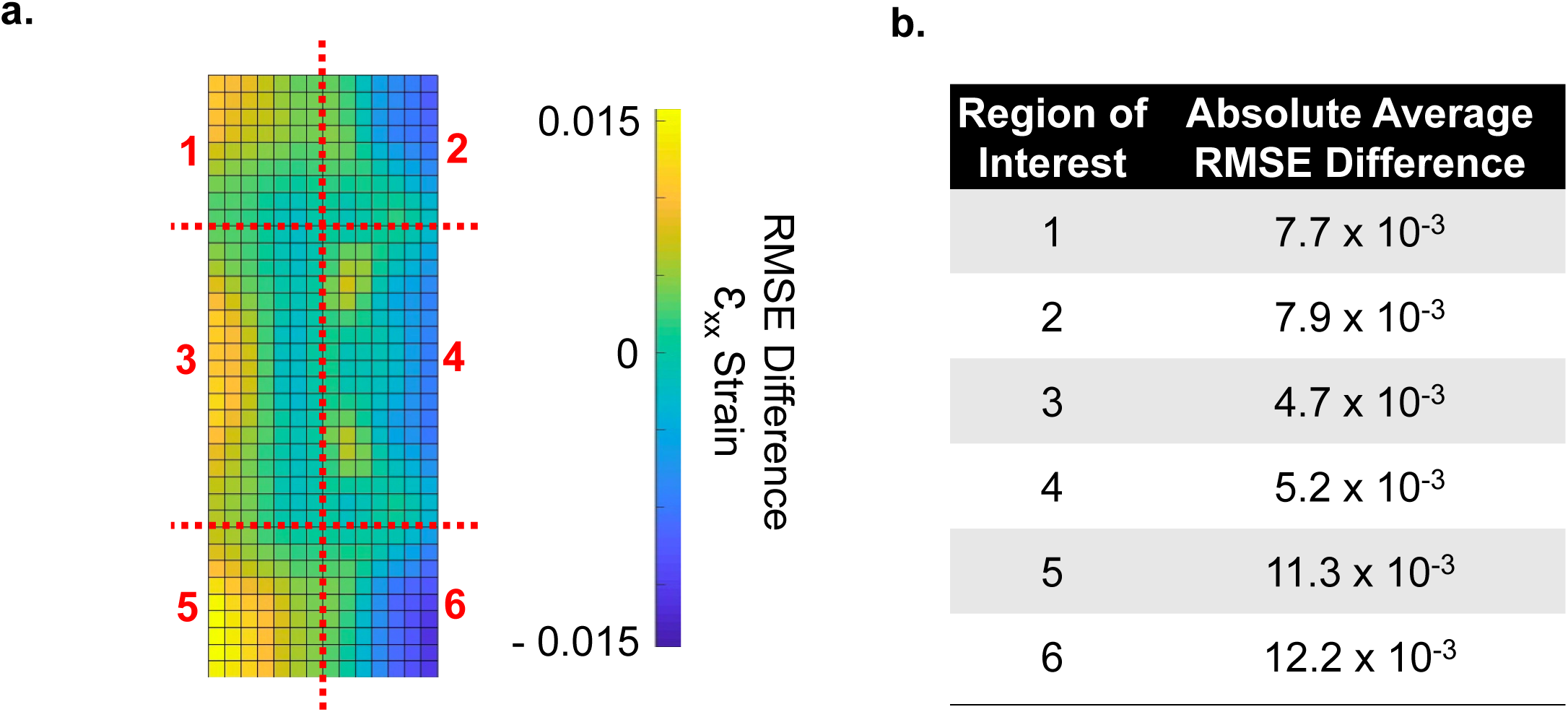
RMSE difference between DIC and FE strains. a) The difference between the surface of DIC experimental and the FE model was measured with a 15×37 grid and divided into 6 regions for comparison. Most of the difference occurs on the outside, especially the outer edges. **b)** Largest difference of 0.015 of averaged ε_xx_ magnitude difference occurs in the regions 5 and 6. Regions 3 and 4 are the closest matching, with an averaged ε_xx_ magnitude difference of 0.0047 and 0.0052, respectively. The corner regions (1, 2, 5 and 6) has an averaged ε_xx_ magnitude difference of 0.0077, 0.0079, 0.0113, and 0.0122, respectively.

**Table 1:** DOE factors used to minimize RMSE between *in vitro* and *in silico* results for calibration gels.

| Stiffness | Friction | Damping | RMSE |
| --- | --- | --- | --- |
| 1.5 | 10 | 0.3 | 0.41 |
| 1.5 | 10 | 0.003 | 0.37 |
| 1.5 | 10 | 0.03 | 0.47 |
| 1.5 | 0.1 | 0.3 | 0.50 |
| 1.5 | 0.1 | 0.003 | 0.38 |
| 1.5 | 0.1 | 0.03 | 0.38 |
| 1.5 | 0.75 | 0.3 | 0.35 |
| 1.5 | 0.75 | 0.003 | 0.36 |
| 1.5 | 0.75 | 0.03 | 0.36 |
| 0.5 | 10 | 0.3 | 0.53 |
| 0.5 | 10 | 0.003 | 0.45 |
| 0.5 | 10 | 0.03 | 0.56 |
| 0.5 | 0.1 | 0.3 | 0.58 |
| 0.5 | 0.1 | 0.003 | 0.50 |
| 0.5 | 0.1 | 0.03 | 0.49 |
| 0.5 | 0.75 | 0.3 | 0.63 |
| 0.5 | 0.75 | 0.003 | 0.64 |
| 0.5 | 0.75 | 0.03 | 0.63 |
| 1 | 10 | 0.3 | 0.29 |
| 1 | 10 | 0.003 | 0.28 |
| 1 | 10 | 0.03 | 0.28 |
| 1 | 0.1 | 0.3 | 0.16 |
| 1 | 0.1 | 0.003 | 0.16 |
| 1 | 0.1 | 0.03 | 0.16 |
| 1 | 0.75 | 0.3 | 0.30 |
| 1 | 0.75 | 0.003 | 0.28 |
| 1 | 0.75 | 0.03 | 0.19 |

### Scaffold finite element modeling

Scaffolds were generated using two bone volumes, 13% and 25%, respectively corresponding to bone volumes of 8 week old (young) or 64 week old (aged) male C57BL/6J mice[66]. Gyroid geometries representing trabecular structures were generated using the equation: *sin*(*x*)*cos*(*y*) + *sin*(*y*)*cos*(*z*) + *sin*(*z*)*cos*(*x*) < *t*, where t is a constant linearly related to the percent volumes[63]. For 13% and 25%, the t scaling value used was –0.121 and –0.752. These models were generated using *MathMod v8.0*, meshed in *Adobe MeshMixer v3.5* using a cell surface density of 128. The models then were scaled to final dimensions of, 5mm x 5mm x 10mm. These then were exported to *Altair Hypermesh v2017.2,* where the STL models were meshed using mixed elements with an element size of 0.14mm. Meshes were made into solid C3D4 elements using tetramesh. The surface of R3D3 elements were exported into ABAQUS.

The FE model was simulated in *ABAQUS (R2019×, Simula, RI).* Pressure-overclosure with a slope of 1.0×10^-2^ was applied to ensure contact between the scaffold and hydrogel surfaces, and both the surface damping and friction coefficients were taken from the optimization procedures. Similar to the previous simulations, the rigid bone was driven by a connector actuator using a periodic velocity equation with an amplitude of 15.61 m/s, and a circular frequency of 100 Hz (628.3 radians). The volumetric strain components corresponding to 13% and 25% scaffolds (ε_xx_, ε_yy_, ε_zz_, ε_xy_, ε_yz_, ε_zx_) were exported. The total hydrogel strain magnitude approximated via equivalent von Mises strain 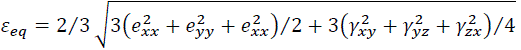, where engineering strains defined as γ*_ij_* = 2ε*_ij_* and deviatoric strains defined as 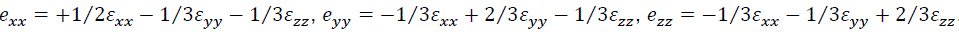.

### Scaffold fabrication and application of low intensity vibration

STL files of the 13 and 25% gyroidal scaffolds were 3D printed using PLA on a PRUSA Mk3s 3D printer (PRI-MK3S-KIT-ORG-PEI) with 0.1mm layer height, and 0.5 x0.5 x1cm overall outer dimensions. Scaffolds were sterilized with ethylene oxide autoclave prior to cellular encapsulation.

HA hydrogels were functionalized with integrin binding motifs (acrylate-PEG-GRGDS) to encourage cell adhesion and metalloproteinases (MMPs)-sensitive crosslinkers (acrylate-PEG-PQ-PEG-acrylate) to allow cells to modify the hydrogel during growth and cell network formation[67]. Cells mixed into the hydrogel at 1×10^6^ cells/ml prior to casting and cross-linking. Cell seeded hydrogels were kept for 7 days in growth media in 5% CO_2_ and 37°C to allow growth and attachment prior to experimentation. After 7 days, scaffolds were transferred to two 6 well plates (Corning#3516) with Growth Media (IMDM, 10% (v/v) FCS and 1% (w/v) pen-strep). Scaffolds were oriented identically to the 3D print, and with the long axis orthogonal to the axis of vibration. Scaffolds were secured with a custom 3D printed PLA top and one 3D printed 30mm PLA platen attached to a stainless-steel screw threaded through the top in each well (**Fig. 7a**). These ensured the scaffolds would not move during vibration. The scaffolds vibrated for one hour each day for 14 days.

### Immunostaining

Samples were rinsed twice in PBS fixed using 4% (v/v) paraformaldehyde for 15 minutes at room temperature. Next, they were rinsed 2 times for 5 mins in PBS before being permeabilized in 0.1% Triton/PBS (v/v) for 15 mins at room temperature. All further steps have a 3 x10 minute rinse in PBS between them. The scaffolds were separated further to be stained against collagen-I (*Bioss Antibodies #bs-10423R-A594)*. Based on antibody species, fixed cells blocked using appropriate serum for 30 minutes at room temperature, stained with primary antibody for 1h at 37° C and labeled against F-actin (1:60 in PBS *Alexa Fluor® 488 Phalloidin-Invitrogen, # A12379*) and DNA (NucBlue, 2 drops/ml, Fisher Scientific #R37605).

After a final two rinses for five minutes each, the scaffolds were imaged in PBS using A1R/MP+ Confocal/Multiphoton Microscope (Nikon Instruments).

### RNA isolation and Assessment of Transcript Levels

Each scaffold was centrifuged at 3000 rpm to separate the gel and scaffolds. Separated gels were added to a single 2 ml tube, and TRIzol™ (*Fisher Scientific#15-596-026*) was added. Gels were mechanically broken up for 30 seconds with a P1000 micropipette and the scaffolds that had been spun to remove the gel were suspended in Zymo lysis buffer and frozen at –80° C overnight. The RNA then was extracted according to manufacturer’s protocols (Zymo Research #R2071), the RNA extracted from cells in the hydrogel and coating the scaffold were combined. These RNA samples then were converted to cDNA and PCR analysis was performed for collagen-I transcripts using 18S as the control gene. Reverse transcription was performed with 1 μg RNA in a total volume of 20 μl per reaction. 25μL amplification reactions contained primers at 0.5 μm, deoxynucleotide triphosphates (0.2 mm each) in PCR buffer, and 0.03 U Taq polymerase along with SYBR-green (Molecular Probes, Inc., Eugene, OR) at 1:150,000. mRNA levels were compared using ΔΔCt method as previously reported.[48] Samples were analyzed in triplicate. PCR products were normalized for the amount of 18S amplicons. The following primers were used for 18S (Forward-GAACGTCTGCCCTATCAACT, Reverse-CCAAGATCCAACTACGAGCT) and collagen-I (Forward-ATGTGCCACTCTGACTGGAA, Reverse-CAGACGGCTGAGTAGGGAA) transcripts.

### Sample Statistics

Results were presented as mean ± standard deviation. Immunostaining and qPCR analyses were performed using samples from three or more independent experiments. As we reported, [13, 49] for comparisons regarding qPCR, differences among treatments within each biological replicate were assumed to follow a normal distribution due to large mean sample size (20,000 or more cells/group), thus for these comparisons, we used two-tailed un-paired one way ANOVA followed by Newman-Keuls post-hoc tests. For other comparisons with smaller sample sizes, including imaging experiments, we used non-parametric two-tailed Mann-Whitney U-test or Kruskal-Wallis test; p-values of less than 0.05 were considered significant. Statistical analyses were performed using Graphpad Prism v.10 (GraphPad Software).

## 3. Results

### Lateral Strain positively correlates with distance during low intensity vibration

High speed visualization of horizontally vibrating wells at 100Hz frequency and 1g acceleration magnitude (**Fig. 3b**) showed that the maximum lateral ε_xx_ strain, measured at the peaks of the cycle (t=π/2 & t=3π/4), reached to 0.014±0.00047 at 8mm spacing over 5 cycles when averaged across three experiments. The maximum ε_xx_ values were symmetrical on both ends of the well, ensuring that steady state was achieved during measurements (**Fig. 3b**). Minimum strains were measured at the end of each positive stroke (t= π). Compared to 8mm at π/2 & t=3π/4 timepoints, averaged maximum ε_xx_ strains of 6mm, 5mm, 4mm well widths were decreased to 0.00806±0.00050 (–0.86%, p=0.66), 0.0075±0.00047 (–7.6%, p=0.17), 0.0030±0.00014 (–62%, p<0.05), respectively (**Fig. 3c**). The π/2 sets for each spacing over all three trials were collated into data distribution plots to show the ε_xx_ strains as a function of distance from the well wall. (**Fig. 3d**).

### Differences between experimental and computational lateral strains are affected by material stiffness and friction and damping coefficients

Prior to validating the FE model, we first optimized the model parameters using the experimental data from the 8mm well width experiments. The effect of material stiffness on the average RSME match between DIC experiments and FE models was quantified using the averaged ε_xx_ distribution at the positive and negative peaks. Using experimentally measured stiffness and FE software default values of 0.75 and 0.03 for the friction and damping coefficients, the average RSME between DIC and FE at π/2 timepoint was 0.19 while the average RMSE between experimental replicates (among the three wells) was 0.15, showing that the average mismatch between FE and DIC was within experimental error. Using default friction and damping coefficients, increasing or decreasing the measured material stiffness by 50% resulted in average RMSE values of 0.36 and 0.63, which were larger than the experimental variation and indicated the importance of accurate stiffness measurements for modeling.

Because the exact boundary conditions between the gels and the PLA were unknown, we investigated the possible combinations of friction and damping coefficient values that could further minimize the average RMSE values. Varying the friction coefficient between 10 and 0.1 resulted in the average RMSE values of 0.28 and 0.16 at measured stiffness and default damping coefficient. Similarly, only varying damping coefficient between values of 0.3 and 0.003 resulted in average RMSE values of 0.28 and 0.3, respectively. Using the experimentally measured mechanical properties, an empirical linear DOE model using a 2^nd^ order polynomial determined the optimal friction / damping combination (**Table 1**). The lowest average RMSE of 0.16 was obtained at the friction/damping combination of 0.1/0.003. Shown by the 15×37 RMSE grid (**Fig. 4a**), maximum differences between the experimental– and FE-derived ε_xx_ magnitudes were largest near the outer walls. Of this, the largest RMSE difference of 0.015 occurred in regions 5 and 6. Match in regions 3 and 4 was closer, an averaged difference of 0.0047 and 0.0052, respectively. Differences in corner regions 1, 2, 5 and 6 were 0.0077, 0.0079, 0.0113, and 0.0122, respectively. RMSE grid for the HA gels is shown in the supplementary **Fig.S3**. The largest RMSE difference of 0.024 again occurs in region 6, with regions 3 and 4 (the middle regions) matching the closest with ε_xx_ averages of .0091 and 0.0082, respectively. Differences in corner regions 1, 2, 4, & 5 were 0.011, 0.010, 0.010, and 0.098, respectively.

Using the friction/damping coefficient combinations that minimized the average RMSE values between DIC and FE results at 8mm well width, we quantified the averaged ε_xx_ magnitudes over 6mm, 5mm, and 4mm for the calibration gel (**Fig. 5a**). The maximum ε_xx_ values were symmetrical at both the π/2 and t=3π/4, in addition to both ends of the well, demonstrating steady state (**Fig. 5b**). The π/2 sets for each spacing over all three trials collated into data distribution plots (**Fig. 5c**), which showed increasing extrema differences as the spacing decreased from 8mm. When compared to 8mm, the averaged maximum ε_xx_ strains of 6mm, 5mm and 4mm well widths decreased to 0.017±0.00040 (–22%, p<0.05), 0.013±0.00030 (41%, p<0.05), 0.010±0.00037 (–54%, p<0.05), and 0.0026±0.00051 (–88%, p<0.05), respectively. Plotting ε_xx_ strains as a function of distance from the well wall showed increasing strain values with larger cavities. (**Fig. 5d**).

**Figure 5.**
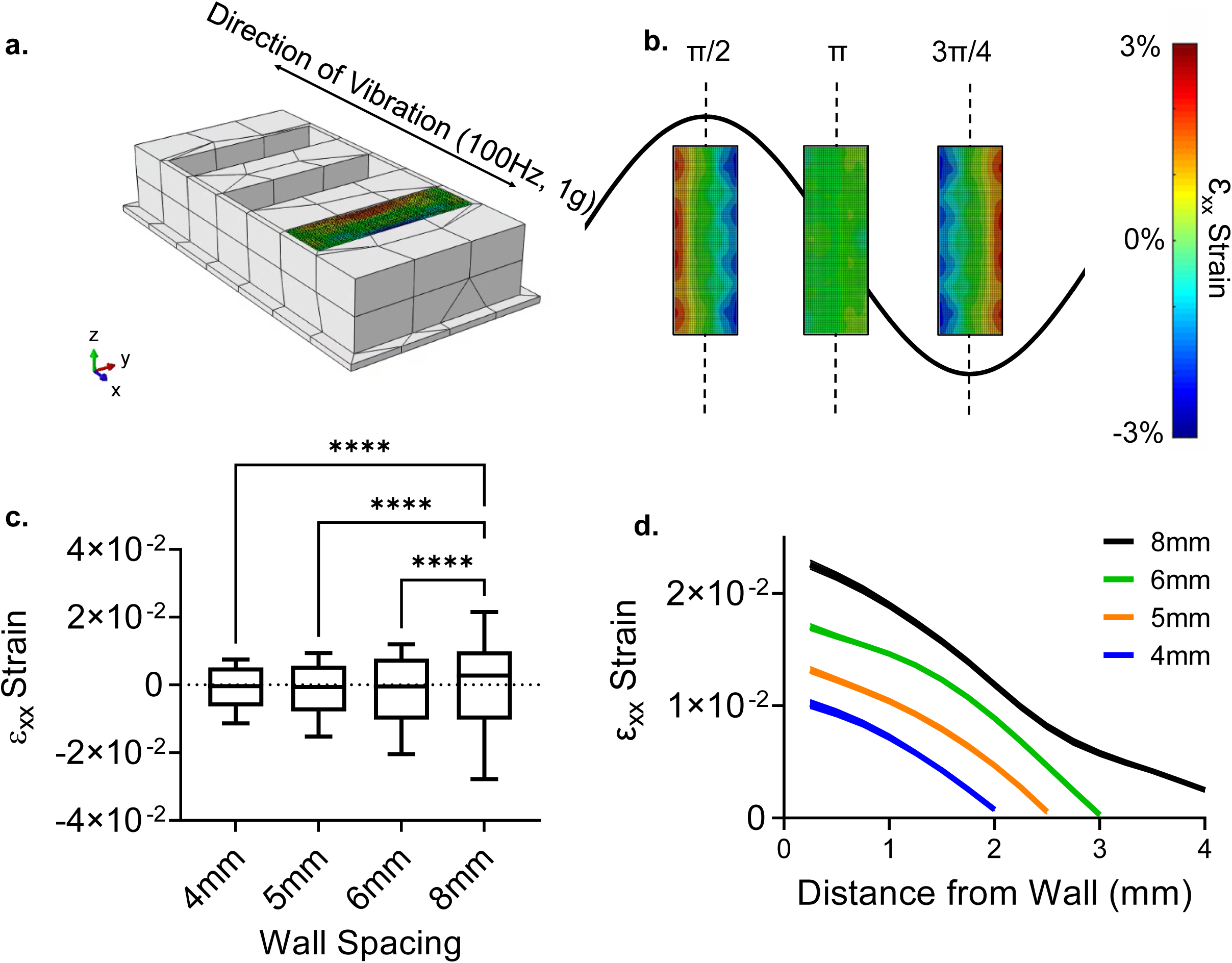
Model validation simulations. a) Finite Element (FE) model setup with 8mm wall spacing. Wells were vibrated identically to the experiments (Fig. 3). Each FE run was proceeded by a gravity step. **b)** ε_xx_ strain magnitude over the cycle. maximum ε_xx_ values were symmetrical at both the π/2 and 3π/4, in addition to both ends of the well, demonstrating steady state. **c)** Extrema differences were evenly spaced as the spacing changes. When compared, the surface strain at the 8mm between the averaged maximum ε_xx_ strains of 6mm,5mm and 4mm well widths decreased to 0.017±0.00040 (–22%, p<0.05), 0.013±0.00030 (41%, p<0.05), 0.010±0.00037 (–54%, p<0.05), and 0.0026±0.00051 (–88%, p<0.05), **d)** Mapping the maximum strain magnitudes, we see a convex parabolic curve with each spacing, excluding 8mm.

### Low trabecular volumes increase LIV-induced hydrogel strains

We next compared the 3D strain distribution between 13% and 25% scaffolds under LIV (**Fig. 6a**) using HA gel mechanical properties. Comparing the Von Mises strain (ε_VM_) magnitude averaged over steady-state cycles showed that ε_VM_ of 25% was 50% smaller compared to 13% scaffold (**Fig. 6b**, p<0.0001). ε_VM_ distribution (**Fig. 6c**) indicated normally distributed strains for both 13% and 25% scaffolds. Histograms of 25% and 13% scaffolds were statistically different (p<0.05), 25% scaffolds had more elements with lower strain values, while 13% scaffolds showed elements with higher ε_VM_ strains. Comparing individual strain components showed a significant difference (p< 0.05) between ε_zz_, ε_xy_ and ε_yz_ strains, no differences were found between ε_xx_, ε_yy_, and ε_zx_ strains. Min/Max values for the normal (εxx, ε_yy_ and ε_zz_) and shear (εxy, ε_yz_ and ε_zx_) components are provided in **Table 2**.

**Figure 6.**
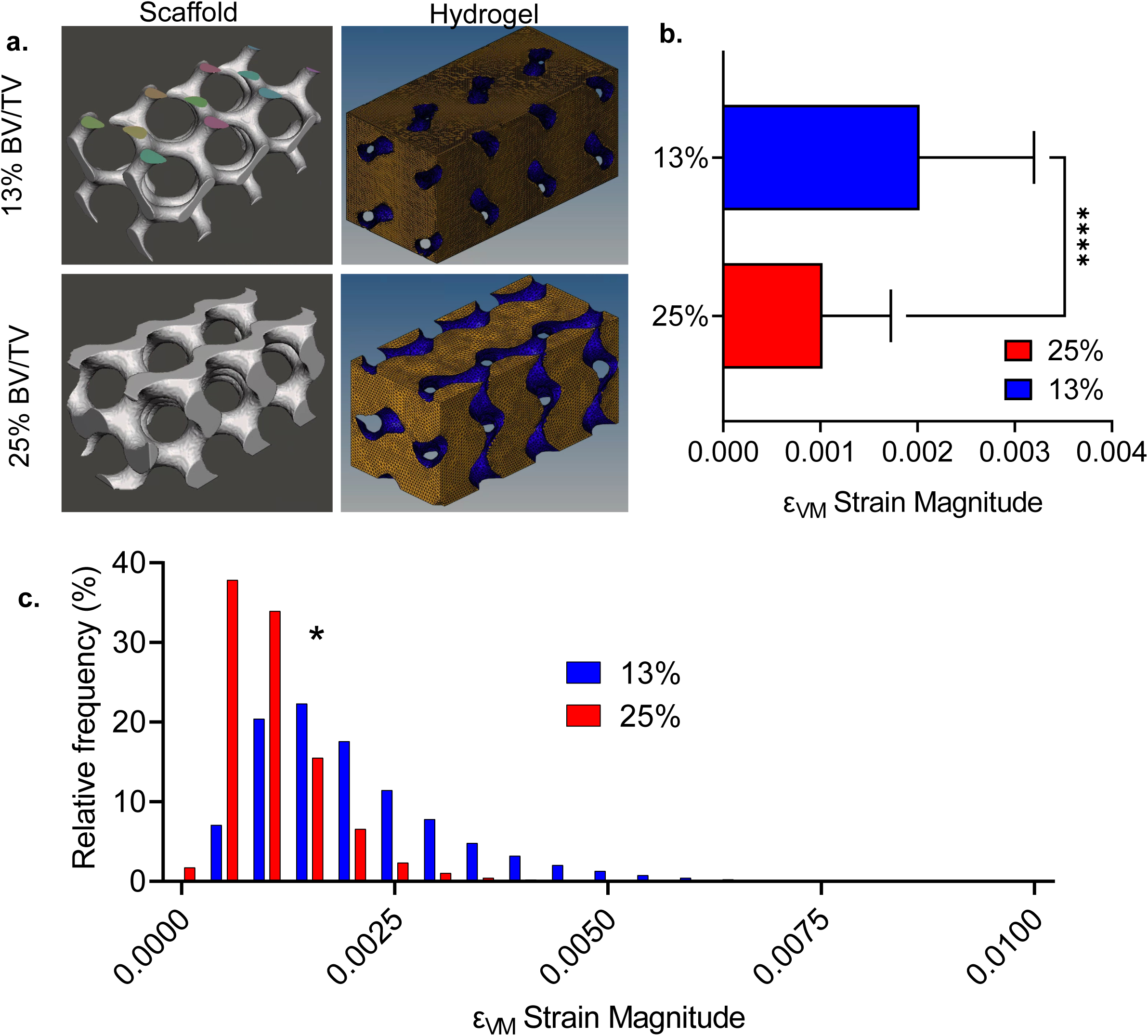
Low trabecular volumes increase LIV-induced hydrogel strains. a) We compared the 3D strain distribution between 13% and 25% scaffolds under LIV using HA gel mechanical properties (Table S1). **b)** Comparing the Von Mises strain (ε_VM_) magnitude averaged over steady-state cycles showed that εVM of 13% was almost doubled compared to 25% scaffold (p<0.0001). **c)** ε_VM_ distribution indicated normally distributed strains for both 13% and 25% scaffolds. Histograms of 25% and 13% scaffolds were statistically different (p<0.05). 25% scaffold had more elements with lower strain values while 13% scaffold showed elements with higher ε_VM_ strains.

**Figure 7.**
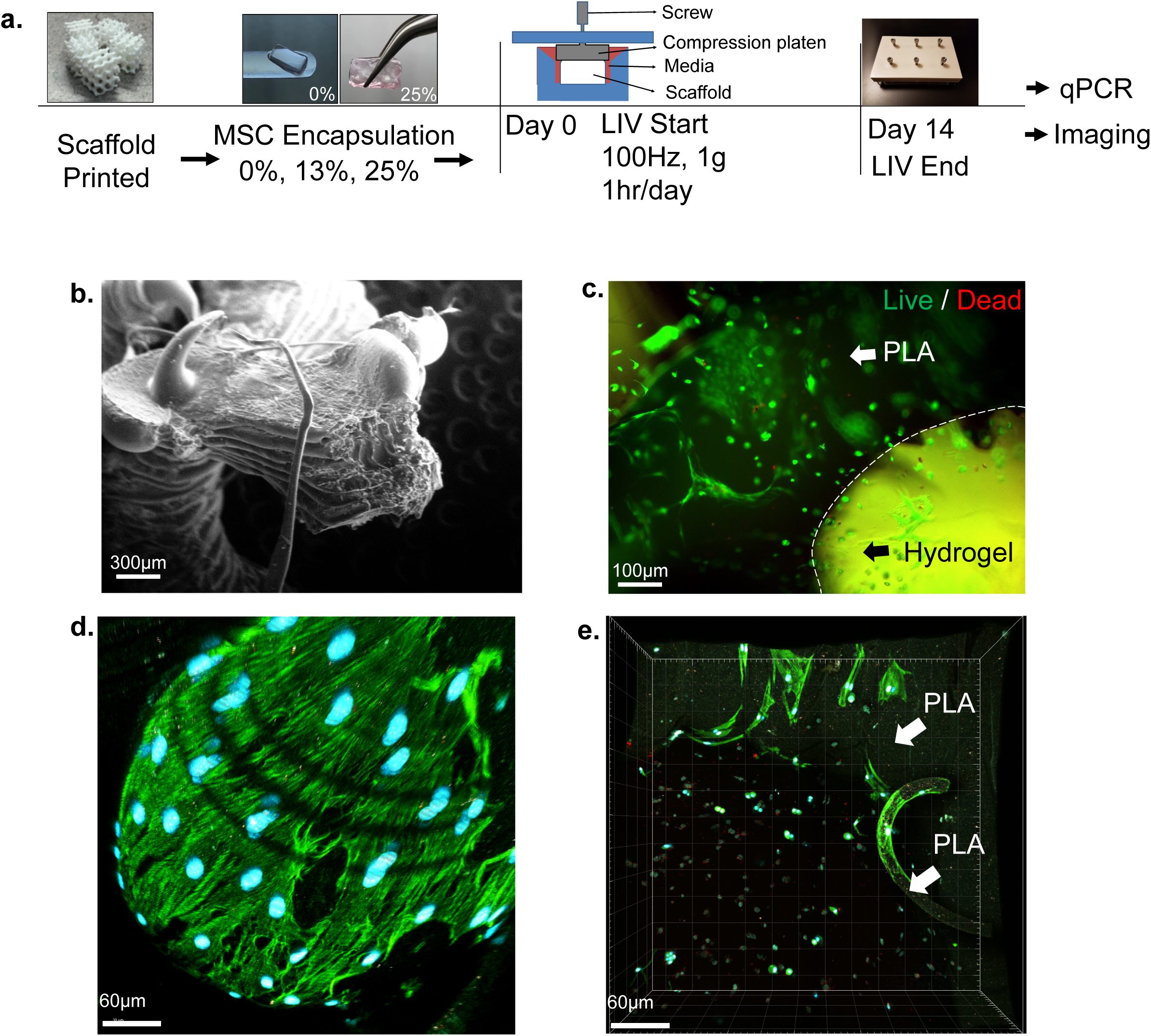
Bone Analogs show viable cells with two distinct cell populations. a) Two sets of PLA scaffolds (13% and 25% volume fill) and 0% controls were generated. Cell-laden 13% and 25% scaffolds were vibrated one hour every day at room temperature using 100 Hz frequency and 1g acceleration magnitude outside of the incubator. After 14 days, samples were allocated for qPCR (N=4 scaffolds/group), imaging (N=3 scaffolds/group) **b)** PLA surface provides an ideal surface for cell attachment **c)** Utilizing live/dead straining to detect cell viability prior to start of the experiments at 7 days post-encapsulation showed viable cells on both PLA and in hydrogels. **d)** Morphology of cells on PLA surfaces was similar to cells on 2D culture plates (Fig.S4), showing elongated morphology while cells in e) hydrogels were smaller and more rounded.

**Table 2:** *In silico* comparison of directional strains between 13% and 25% scaffolds during LIV.

| Tensor | 13% |  | 25% |  |
| --- | --- | --- | --- | --- |
|  | Min | Max | Min | Max |
| XX | -0.053 | 0.062 | -0.047 | 0.012 |
| YY | -0.056 | 0.071 | -0.017 | 0.050 |
| ZZ | -0.075 | 0.067 | -0.047 | 0.029 |
| XY | -0.102 | 0.067 | -0.023 | 0.025 |
| YZ | -0.082 | 0.104 | -0.042 | 0.053 |
| ZX | -0.078 | 0.089 | -0.021 | 0.062 |

### Bone Analogs show viable cells with two distinct cell populations

In order to compare the possible differences in MSC mechanoregulation due to bone volume differences we generated two sets of PLA scaffolds (13% and 25% volume fill) and 0% controls. Cell-laden 13% and 25% scaffolds vibrated one hour every day at room temperature using 100Hz frequency or 1g acceleration magnitude (**Fig. 7a**). During vibrations, controls were kept outside of the incubator but were not vibrated. After 14 days, samples were allocated for qPCR (N=4 scaffolds/group), imaging (N=3 scaffolds/group). Live/dead straining prior to start of the experiments at day 0 (7 days post-encapsulation) showed viable cells on both PLA and in hydrogels (**Fig. 7c**). Shown in **Fig. 7d**, morphology of cells on PLA surfaces were similar to 2D culture plates (**Fig.S4**), showing elongated morphology, while cells in hydrogels were smaller and more rounded (**Fig. 7e**).

### LIV increases F-actin volume independent of hydrogel strains

We first tested the F-actin volume in hydrogels in response to LIV using samples stained against Phalloidin. Comparing 0%, 13% and 25% samples without LIV showed no significant change in the F-actin volume. Shown in **Fig. 8b**, comparing –LIV samples with their +LIV counterparts we found that 13% BV/TV group with a larger hydrogel strain (average ε_VM_ 0.2%, **Fig. 6b**) had a trend toward increasing F-actin (+50%, ns) while 25% BV/TV with a lower hydrogel strain (average ε_VM_ 0.1%, **Fig. 6b**) showed a 2.4-fold increase (p<0.05), suggesting that LIV effect is independent of the hydrogel strain.

### Trabecular volume differentially regulates basal and LIV-induced collagen-I production

Collagen-I immunostaining showed a more robust collagen labeling in vibrated samples (+LIV), when compared to non-vibrated controls (–LIV, **Fig. 8a-b**). Collagen-I mRNA quantification showed a trend of increased collagen-I levels in +LIV samples (+50% in 13% and +94% in 25% samples, **Fig. 5c**). Quantification of collagen-I from volumetric scans showed unchanged basal collagen-I levels between 0% and 13% controls while –LIV 25% group had a 73% increase compared to –LIV 13% group (p<0.05). LIV increased collagen-I in both 13% and 25% groups by 87% and 53% respectively (p<0.01) and +LIV 25% remained significantly higher compared to +LIV 13% group (37%, p<0.05).

## 4. Discussion

We found that hydrogel strain magnitudes between scaffolds representing young (25%) and aged (13%) trabecular volumes significantly differ during LIV. Validation studies indicate that strain magnitude within the LIV direction is positively correlated with the spacing between two solid structures (**Fig. 3d & 5c**). This conclusion was supported when comparing 25% and 13% scaffolds: Average ε_VM_ for the 25% scaffold was 50% smaller when compared to the 13% scaffold which had more spacing between trabecular struts (**Fig. 6a-b**). These findings indicate that age-related bone adaptations, decreased trabecular volume and increased cortical circumference[56] may serve to increase the magnitude of bone marrow deformation during physical activity.

In contrast to larger hydrogel strains, observed F-actin volumes (**Fig. 8**) and collagen-I production (**Fig. 9**) of 13% scaffolds were lower when compared to 25% scaffolds. Given that LIV-induced outcomes were lower in 13% scaffolds, we conclude that LIV response is relatively insensitive to hydrogel strains. This insensitivity of LIV response in hydrogels can be attributed to several factors. First, measured strain differences within the hydrogel only represent a fraction of the mechanical information. During 1g, 100Hz sinusoidal vibrations the entire scaffold, including PLA surfaces and hydrogel, accelerates and decelerates to speed of ±15mm/s, 100 times each second, providing another mechanical input in addition to LIV-induced hydrogel strains. We previously reported that when substrate strains are extremely low, LIV activates COX-2 signaling in MC3T3 osteoblasts[54], improves gap junctional communication in MLO-Y4 osteocytes[55] and increases proliferation as well as biosynthesis of collagen and related proteins in MSCs under simulated microgravity[13]. While this suggests that LIV could regulate collagen production and changes in F-actin that are independent of hydrogel strains, the finding that 25% scaffolds produce more collagen when compared to 13% scaffolds and 0% controls indicates that loss of bone volume could negatively affect the function of the MSCs inside the hydrogel.

**Figure 8.**
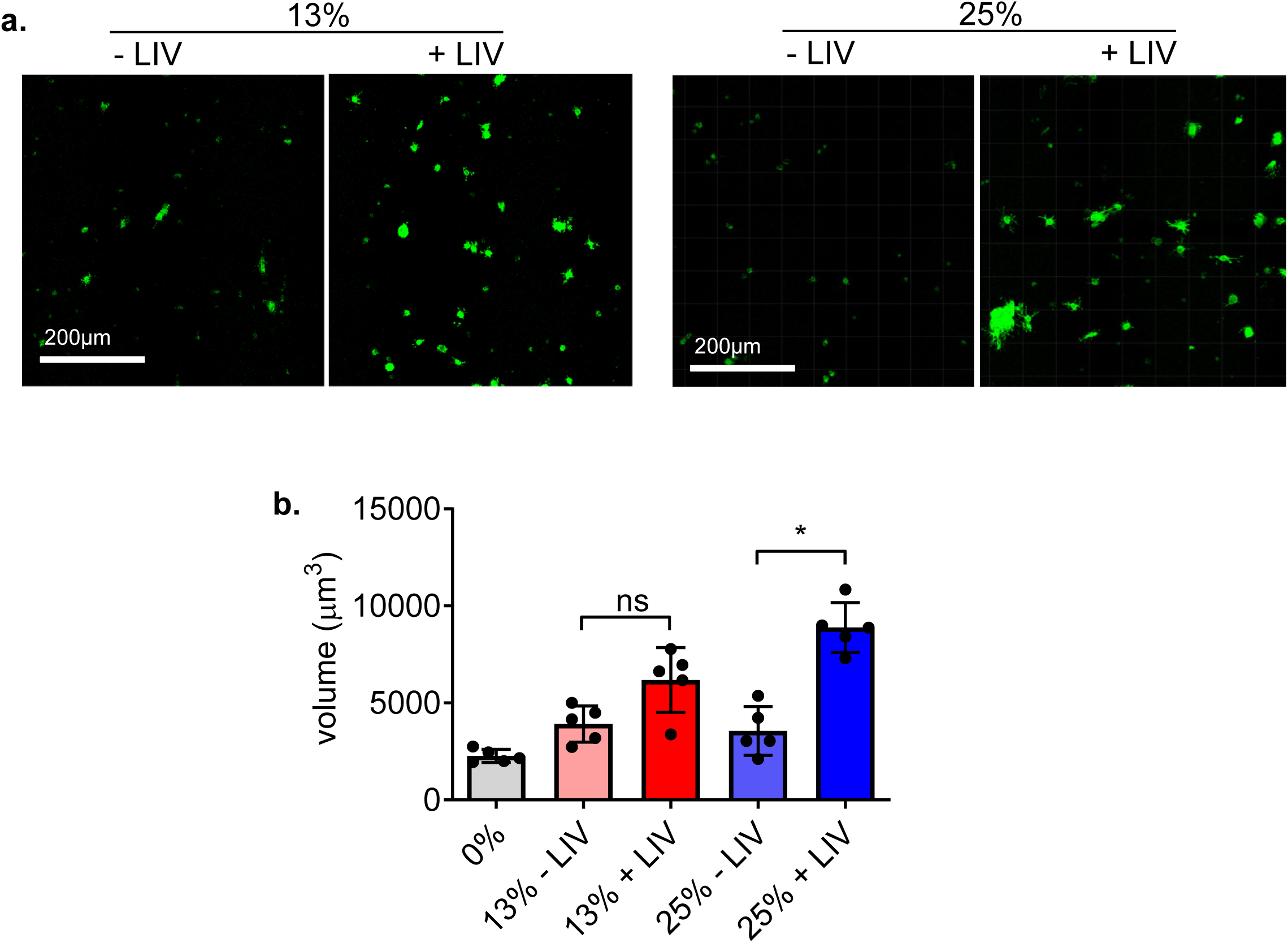
LIV increases F-actin volume. F-actin staining in **a)** cells in 13% and 25% scaffolds show increased F-actin volume in +LIV samples when compared to controls (–LIV). **b)** Quantification F-actin volume showed no change in –LIV groups. 13% +LIV groups trended toward increased F-actin and 25% +LIV group showed 2.5-fold increase in F-actin volume (p<0.05).

**Figure 9.**
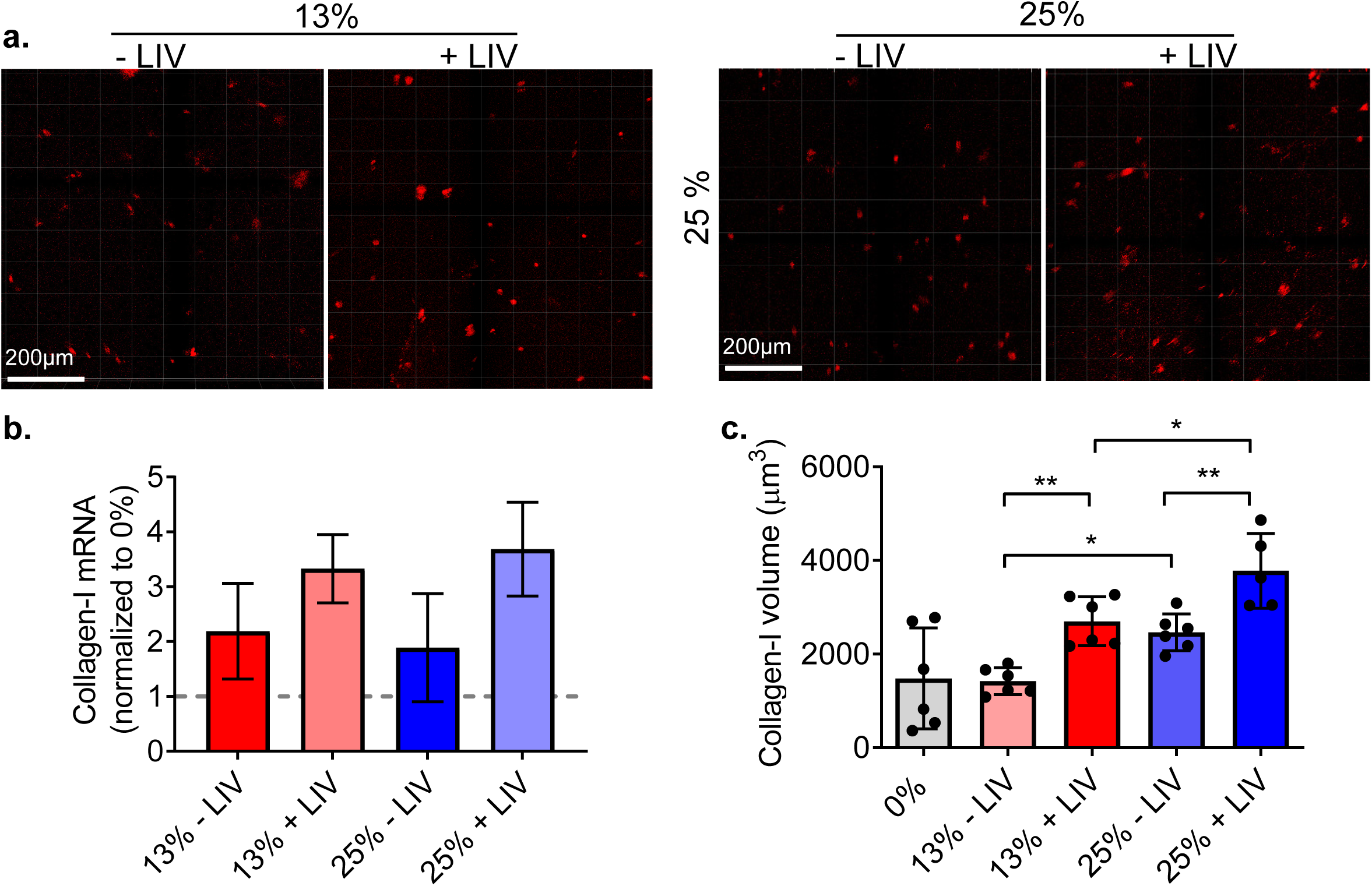
Trabecular volume differentially regulates basal and LIV-induced collagen-I production. Collagen-I immunostaining in **a)**13% and 25% scaffolds show increased collagen labeling in vibrated samples (+LIV), vs non vibrated controls (–LIV). **b)** Collagen-I mRNA quantification show an increasing trend of collagen-I levels in +LIV samples (+50% in 13% and +94% in 25% samples. All samples were normalized to 0% controls (dashed gray bar) **c)** Quantification of collagen-I volume showed unchanged basal collagen-I levels between 0% and 13% controls while –LIV 25% group showed a 73% increase compared to –LIV 13% group (p<0.05). LIV increased collagen-I in both 13% and 25% groups by 87% and 53% respectively (p<0.01) and +LIV 25% remained significantly higher compared to +LIV 13% group (37%, p<0.05).

A possible reason for increased collagen production is the difference in scaffold surface area. Due to increased scaffold volume, the 25% scaffolds had a smaller hydrogel volume, increasing the possibility that imaging regions are closer to the scaffold walls. Confocal images showed many cells that were viable and spread on the PLA surfaces (**Fig. 7**). While our analysis excluded cells on the PLA surface from the data, these cells on PLA surfaces are likely to produce paracrine signals to cells within the hydrogels. For example, during bone remodeling, early stage osteoblast lineage cells that reside on bone surfaces are at a low cell density, but their density increases during the reversal phase [68]. Interestingly, early osteoblast lineage cells are “pro-resorptive” and osteoid deposition starts only when a critical cell density is reached [69]. This suggests that basal and LIV-driven density changes of these PLA surface cells could not only signal to hydrogels cells, but also could regulate important factors such YAP/TAZ that change their subcellular localization based on cell density[70]. While the metrics of this population were not investigated in this study, future studies aimed at possible interactions of cells on the PLA surface and hydrogel will be critical to identify the interactions between scaffold adherent and non-adherent cell populations.

Despite our attempts to calibrate material parameters in the FE model based on experimental data, differences between our DIC experimental data and our FE model data were seen in both calibration and HA gels. Highlighting the inherent variation in DIC experiments, the average RMSE values between calibration or HA experiment were close to differences between experimental data and computational models. Several factors could have contributed to this variability. The first relates to the rigid body assumption we assigned to PLA in our simulations. While this appears to be a reasonable assumption and used by others[57], a potential model limitation is the inability to account for strain damping from the walls, which could decrease the wall strain in the FE model. Another source of error is the DIC method itself, where the PLA-hydrogel boundaries cannot be resolved via optical resolution, resulting in lower averages at the boundaries when compared to FE model, where resolution is numerical. While, this limitation of DIC could underestimate the magnitudes towards the walls, FE model did predict the relative differences observed experimentally among different well sizes, and therefore the relative differences predicted by the model between the two bone volume fractions remains credible.

In this study, we developed bi-phasic bone analogs with a fully characterized hydrogel strain environment during LIV. Our findings showed that trabecular densities associated with advanced age resulted in higher hydrogel strains, while trabecular densities expected at adulthood resulted in smaller strains during LIV. Despite the lower strain magnitudes, hydrogels mimicking young bone marrow supported higher F-actin measures and collagen production in both basal and mechanically stimulated scaffolds, indicating that factors other than hydrogel environment are in play. These findings not only reveal interesting new avenues of research to study interactions between scaffold-lining and hydrogel encapsulated cells to mimic bone marrow mechanobiology, but the model itself establishes a versatile 3D platform to study how mechanical signals may affect multicellular processes such as environmental factors, aging, inactivity, microgravity and radiation.

## Data availability

The datasets generated and/or analyzed during the current study are available from the corresponding author on reasonable request.

## Acknowledgements

Awards from NIH R01AG059923, P20GM109095, 5P2CHD086843-03 and NSF 1929188, 2025505, supported this study.

## Author Contributions

Alexander M. Regner: concept/design, data analysis/interpretation, manuscript writing Maximilien DeLeon: concept/design, data analysis/interpretation, manuscript writing Kalin D. Gibbons: concept/design, manuscript writing, final approval of manuscript Sean Howard: concept/design, data analysis, manuscript writing, final approval of manuscript Derek Q. Nesbitt: data analysis, final approval of manuscript Trevor J. Lujan: data analysis, final approval of manuscript Clare K. Fitzpatrick manuscript writing, data analysis, financial support, final approval of manuscript Mary C. Farach-Carson: manuscript writing, data interpretation, financial support, final approval of manuscript Danielle Wu: manuscript writing, data analysis, financial support, final approval of manuscript Gunes Uzer: concept/design, data analysis/interpretation, financial support, manuscript writing, final approval of manuscript

**Table S1:** DOE factors used to minimize RMSE between *in vitro* and *in silico* results for HA gels.

| Friction | Damping | RMSE |
| --- | --- | --- |
| 10 | 0.3 | 0.28 |
| 10 | 0.003 | 0.29 |
| 10 | 0.03 | 0.28 |
| 0.1 | 0.3 | 0.38 |
| 0.1 | 0.003 | 0.25 |
| 0.1 | 0.03 | 0.35 |
| 0.75 | 0.3 | 0.28 |
| 0.75 | 0.003 | 0.28 |

## Supplementary Figures

**Figure S1.**
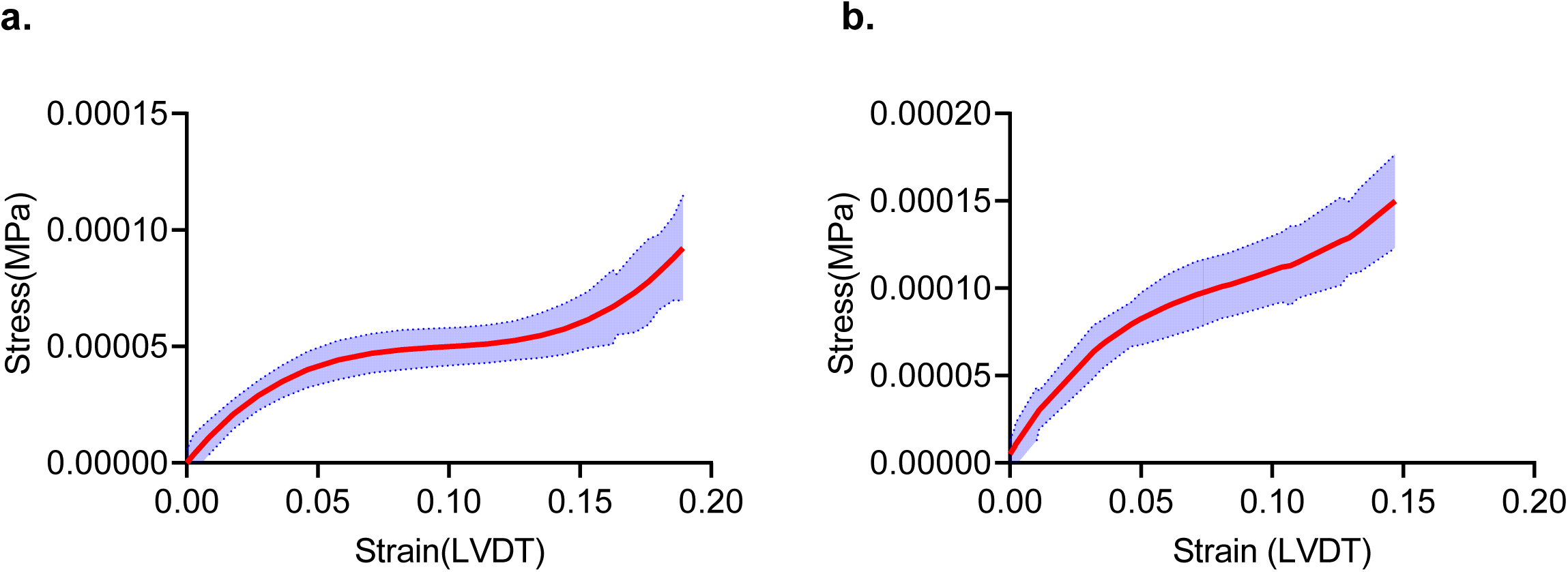
Stress-strain curves for the **a)** calibration and **b)** HA gels from instantaneous compression tests.

**Figure S2.**
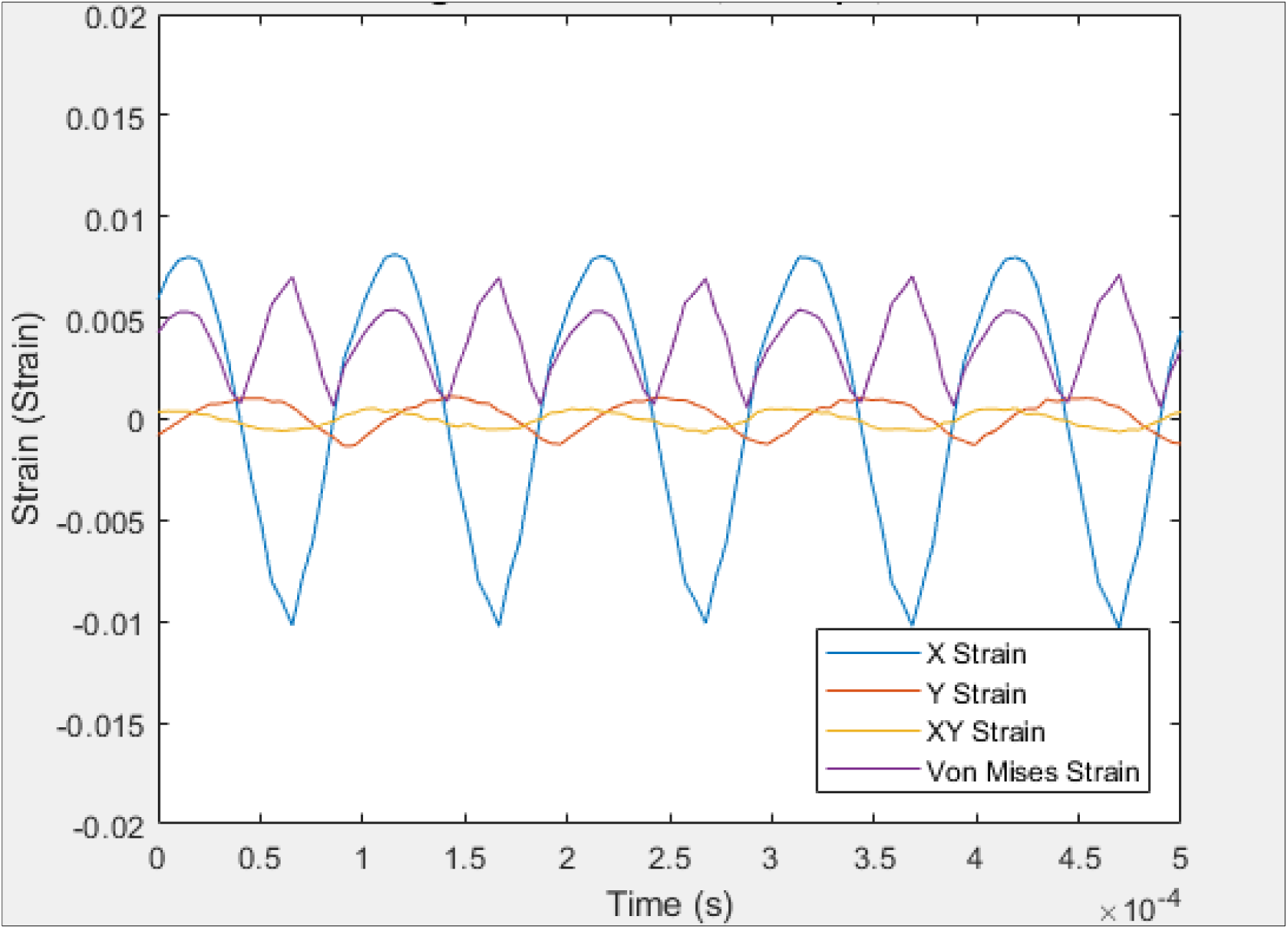
Calibration gel strains were compared during horizontally applied LIV stimulation along x-direction. The ε_xx_ strain was found to be at least one order of magnitude larger than ε_yy_ or ^ε^xy.

**Figure S3.**
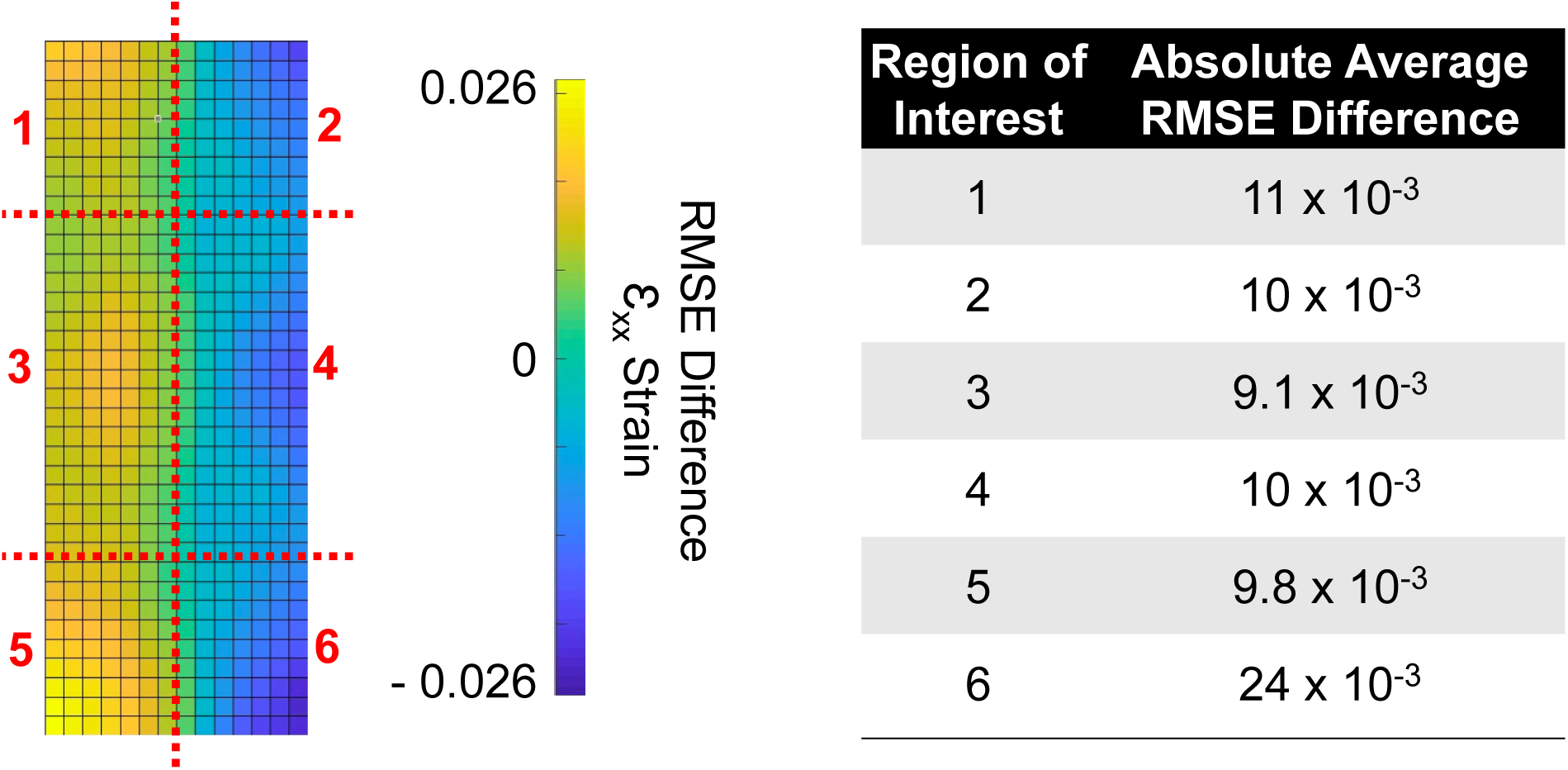
**a**) The difference between the surface of DIC experimental and the FE model was measured with a 15×37 grid and divided into 6 regions for comparison. Most of the difference occurs on the outside, especially the outer edges. **b)** The largest difference of 0.024 again occurs in region 6, with regions 3 and 4 matching the closest with ε_xx_ averages of .0091 and 0.0082, respectively. The corner regions had average differences of 0.011, 0.010, 0.010, and 0.098 for 1, 2, 4, & 5 respectively.

**Figure S4.**
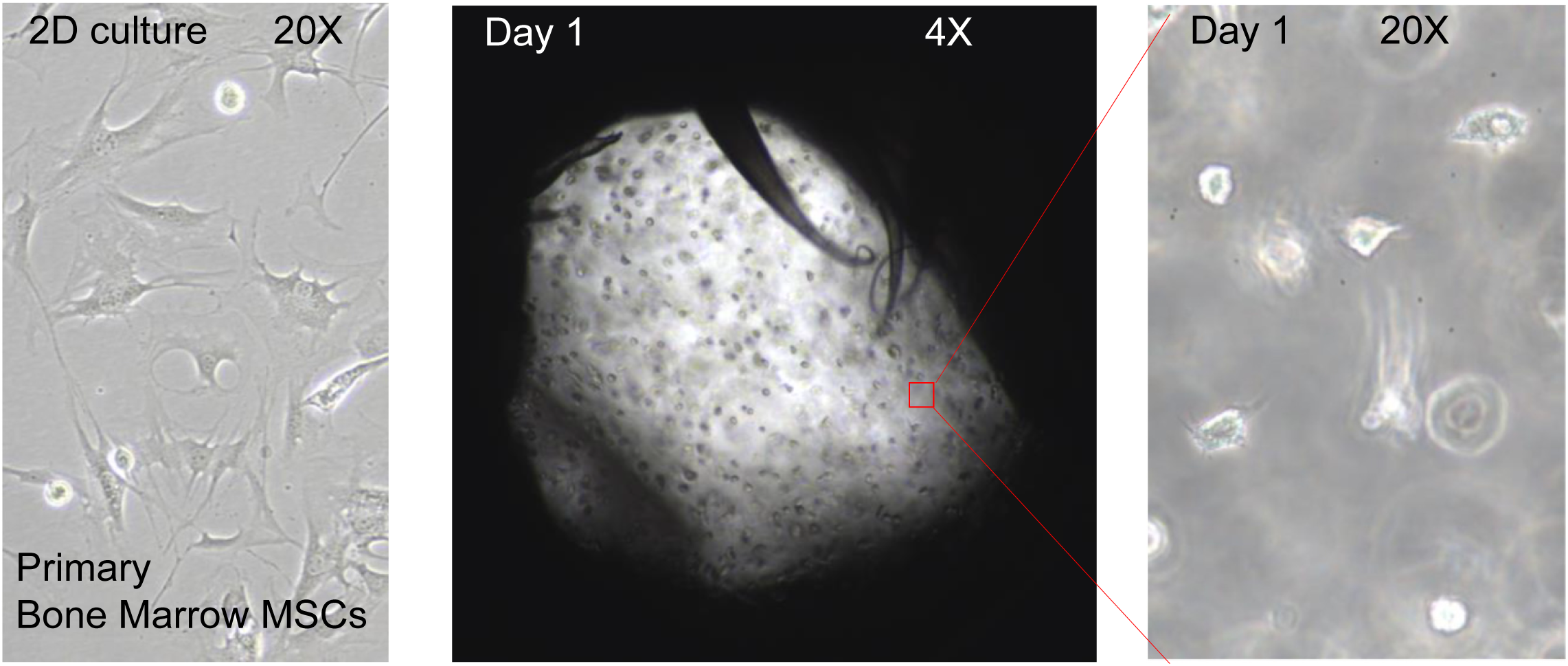
Prior to encapsulation MSCs show elongated morphology on 2D culture plastic. 24hr after the cell encapsulation, MSCs are largely retained within the hydrogel (4X) and show some elongation, suggesting active migration (20×).

